# Inflammation-Induced Claudin-2 Upregulation Limits Pancreatitis Progression by Enhancing Tight Junction-Controlled Pancreatic Ductal Transport

**DOI:** 10.1101/2023.09.01.555960

**Authors:** Sneha Kesaraju, Yueying Li, Junjie Xing, Melissa Tracy, Kristin Wannemo, Aysha Holder, Piao Zhao, Mohammed Azizuddin Khan, Joseph Kainov, Niyati Rana, Mohammed Sidahmed, Sanjiv Hyoju, Lexi Smith, Jonathan Matthews, Savas Tay, Fatemeh Khalili-Araghi, Maunak Rana, Scott A. Oakes, Le Shen, Christopher R. Weber

**Author notes:** These authors contributed equally (Sneha Kesaraju, Yueying Li).

## Abstract

Pancreatitis is an inflammatory disease of the pancreas that can arise due to various factors, including environmental risks such as diet, alcohol, and smoking, as well as genetic predispositions. In some cases, pancreatitis may progress and become chronic, leading to irreversible damage and impaired pancreatic function. Genome-wide association studies (GWAS) have identified polymorphisms at the X-linked *CLDN2* locus as risk factors for both sporadic and alcohol-related chronic pancreatitis. *CLDN2* encodes claudin-2 (CLDN2), a paracellular cation-selective channel localized at tight junctions and expressed in the pancreas and other secretory organs. However, whether and how CLDN2 may modify pancreatitis susceptibility remains poorly understood. We aimed to clarify the potential role of CLDN2 in the onset and progression of pancreatitis.

We employed multiple methodologies to examine the role of CLDN2 in human pancreatic tissue, caerulein-induced experimental pancreatitis mouse model, and pancreatic ductal epithelial organoids. In both human chronic pancreatitis tissues and caerulein-induced experimental pancreatitis, CLDN2 protein was significantly upregulated in pancreatic ductal epithelial cells. Our studies using pancreatic ductal epithelial organoids and mice demonstrated the inflammatory cytokine IFNγ upregulates claudin-2 expression at both RNA and protein levels. Following caerulein treatment, *Ifng* KO mice had diminished upregulation of CLDN2 relative to WT mice, indicating that caerulein-induced claudin-2 expression is partially driven by IFNγ. Functionally, *Cldn2* knockout mice developed more severe caerulein-induced experimental pancreatitis, indicating CLDN2 plays a protective role in pancreatitis development. Pancreatic ductal epithelial organoid-based studies demonstrated that CLDN2 is critical for sodium-dependent water transport and necessary for cAMP-driven, CFTR-dependent fluid secretion. These findings suggest that functional crosstalk between CLDN2 and CFTR is essential for fluid transport in pancreatic ductal epithelium, which may protect against pancreatitis by adjusting pancreatic ductal secretion to prevent worsening autodigestion and inflammation.

In conclusion, our studies suggest CLDN2 upregulation during pancreatitis may play a protective role in limiting disease development, and decreased CLDN2 function may increase pancreatitis severity. These results point to the possibility of modulating pancreatic ductal CLDN2 function as an approach for therapeutic intervention of pancreatitis.

## INTRODUCTION

Pancreatitis is characterized by inflammation and auto-digestion of the pancreas, which sometimes progresses to permanent scarring of pancreatic tissue (Beyer *et al*., 2020). The condition presents with symptoms such as abdominal pain, nausea, vomiting, and steatorrhea. In more severe or recurrent cases, pancreatitis can lead to complications such as malnutrition due to exocrine pancreatic insufficiency, diabetes due to endocrine dysfunction, and an increased risk of pancreatic ductal adenocarcinoma. In the United States, pancreatitis, both acute and chronic forms, affects a significant number of adults, with the annual incidence of chronic pancreatitis alone ranging from 5 to 8 per 100,000 adults, and prevalence estimates between 42 to 73 per 100,000 adults (Vege & Chari, 2022).

While lifestyle factors such as alcohol and tobacco abuse are well-recognized contributors to pancreatitis, genetic predispositions also play an important role in its development. Mutations in genes such as PRSS1, SPINK1, CTRC, CASR, and CFTR have been linked to increased susceptibility to the disease (Singh et al., 2019). Genome-wide association studies (GWAS) have identified the *rs12688220* locus, which is close to the CLDN2 and MORC4 genes, as a risk locus for chronic pancreatitis in North American patients of European ancestry (Whitcomb et al., 2012). Although subsequent analysis has assigned *rs12688220* to *MORC4* gene, additional analyses has identified multiple SNPs, including *rs4409525, rs12008279, rs7057398* within the *CLDN2* locus that are associated with increased risk of pancreatitis (Avanthi et al., 2015; Deng & Li, 2020; Derikx et al., 2015; Giri et al., 2016; Masamune et al., 2015; Wolthers et al., 2017). Despite such genetic evidence, how these SNPs may impact pancreatitis development remains unclear, as none of these SNPs are located within the coding region of the *CLDN2* gene. Furthermore, these studies do not directly address the physiological and pathophysiological roles that CLDN2 may have during pancreatitis development.

Claudins are a 27-member family of tight junction proteins that regulate paracellular permeabilities of small molecules, particularly ions and water. While some claudins form barriers to limit small molecule transport, others create size and charge selective pores that allow specific molecules to pass through the tight junction, thus controlling paracellular transport (Shen *et al*., 2011). Expression of distinctive combinations of claudin family members is thought to contribute to unique transport properties of different epithelia. It has been shown that CLDN2 monomers polymerize within a cell membrane to form strands (Furuse, Sasaki, et al., 1998). Trans-interactions of these strands between cells allow the formation of cation selective ion channels. It has been suggested that claudin channels are formed from four monomers and CLDN2 channels control paracellular Na^+^, Ca^2+^, and water transport (Amasheh et al., 2002; Rosenthal et al., 2017; Samanta et al., 2018; Yu et al., 2010). The increase in CLDN2 expression under inflammation is well established, particularly in diseases of the gastrointestinal tract such as inflammatory bowel disease, infectious colitis, and celiac disease (Denizot *et al*., 2012; Heller *et al*., 2005; Liu *et al*., 2013; Prasad *et al*., 2005; Schmitz *et al*., 1999; Szakál *et al*., 2010). Cytokines, including IFN-γ, IL-6, TNFα, IL-13, and IL-22, are known to modulate the expression of CLDN2 in response to tissue inflammation in a number of different cell types, contributing to changes in paracellular transport and barrier function (Amoozadeh *et al*., 2015; Mankertz *et al*., 2009; Suzuki *et al*., 2011; Tian *et al*., 2018; Wang *et al*., 2017; Weber *et al*., 2010; Yamamoto *et al*., 2004). Given the diverse roles of claudins in regulating paracellular transport, the role of CLDN2 in pancreatic ductal epithelial cells, particularly under inflammatory conditions such as pancreatitis, warrants further investigation.

We demonstrate that CLDN2 expression is upregulated in pancreatic ductal epithelium of chronic pancreatitis patients, a phenomenon also observed in the caerulein mouse model of experimental pancreatitis. Our results show that IFNγ contributes to the upregulation of CLDN2 in pancreatic ductal epithelial cells. Studies using *Cldn2* knockout (KO) mice reveal increased susceptibility to caerulein-induced pancreatic injury, and *Cldn2* KO diminishes cAMP driven / CFTR-dependent fluid transport across pancreatic ductal epithelium. Taken together, our data show that increased CLDN2 expression limits pancreatitis development through regulating paracellular fluid transport in pancreatic ducts.

## RESULTS

### CLDN2 expression is upregulated in human chronic pancreatitis

Previous studies on the pancreas demonstrated that CLDN2 is predominantly expressed in pancreatic ductal epithelial cells, with low expression in pancreatic acinar cells. However, how CLDN2 expression may be altered in chronic pancreatitis remains to be determined. To investigate this, we evaluated CLDN2 expression in pancreatectomy specimens from patients with chronic pancreatitis. Hematoxylin and Eosin (H&E) staining showed acinar cell loss and fibrosis in the chronic pancreatitis samples, which was not present in normal pancreas samples (Fig. 1A). CLDN2 expression was significantly higher in the ducts of chronic pancreatitis tissues compared to normal pancreas, as determined by immunohistochemical staining (n=13 for pancreatitis samples and n=6 for normal pancreas samples; p < 0.0001). (Table 1, Fig. 1B – 1C). Similar results were observed with immunofluorescence microscopy (Fig. 1D). These data suggest CLDN2 may be involved in the pathophysiological processes associated with chronic pancreatitis. In order to define whether CLDN2 upregulation is a specific process or a general alteration of the tight junctions, we also assessed the expression of CLDN1 (Furuse, Fujita, et al., 1998; Inai et al., 1999) and occludin (Buschmann et al., 2013; Furuse et al., 1993), two other important tight junction proteins known to be expressed in pancreatic ductal epithelial cells (Kojima et al., 2013) by immunohistochemistry (IHC) and immunofluorescence, respectively. CLDN1 is a prototypical “sealing claudin,” while occludin plays a critical role in regulating barrier permeability to large molecules. The expression of these proteins remained unchanged, suggesting that CLDN2 upregulation is specific to chronic pancreatitis (Supplementary Figure 1).

**Figure 1.**
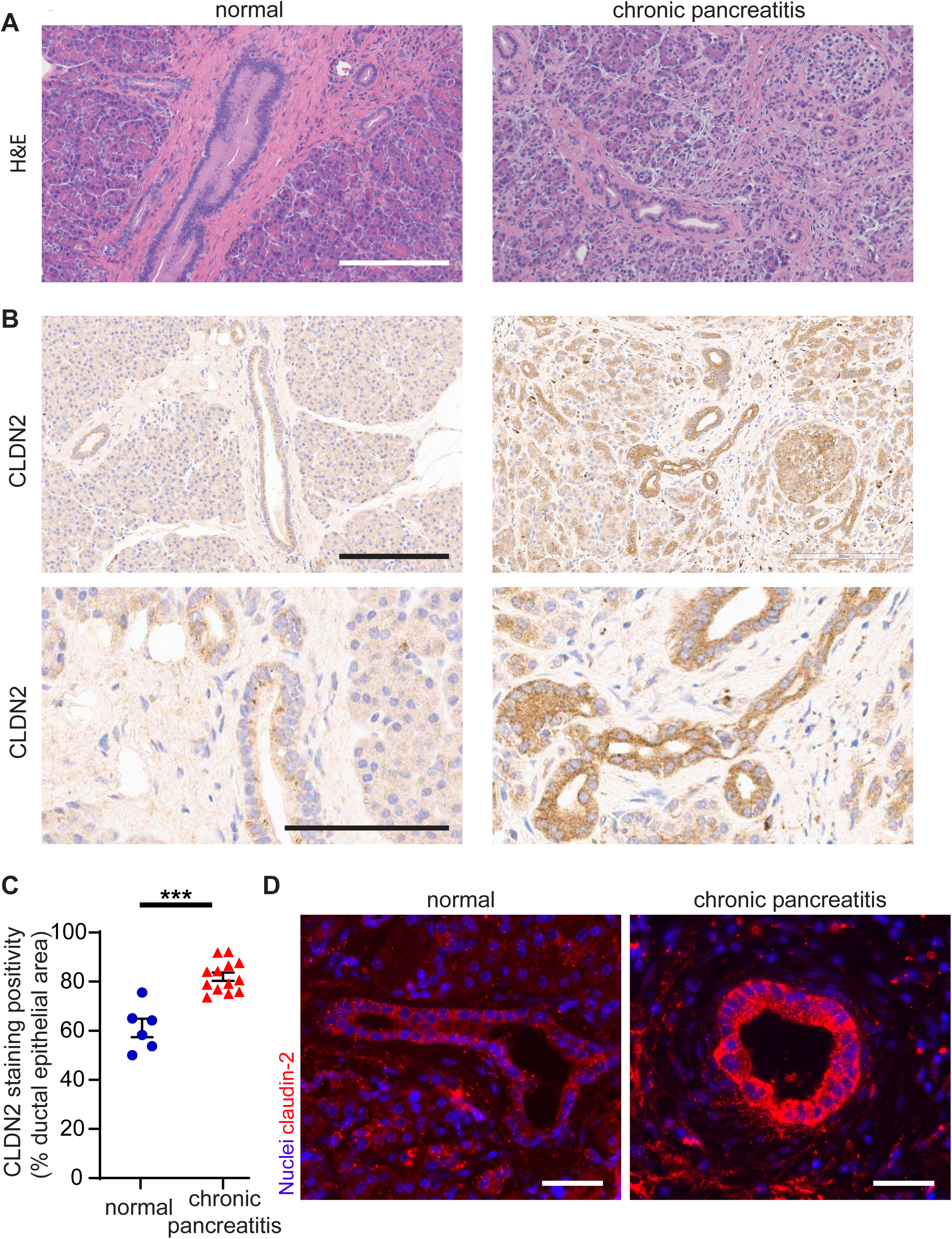
CLDN2 protein expression is upregulated in human chronic pancreatitis. **A.** Representative images of H&E-stained human pancreatic tissue sections. (Scale bar 200 µm) **B.** CLDN2 immunohistochemical staining of human pancreatic tissue sections. (Top scale bar 200 µm, bottom scale bar 100 µm) C. Quantification of pancreatic ductal epithelial CLDN2 staining. Blue circles: normal pancreas tissue (n=6), red triangles: human chronic pancreatitis tissue (n=13) (p < 0.0001). Unpaired Student’s *t* test, ***p<0.001. **D.** Representative immunofluorescence staining of CLDN2 in human pancreatic tissue sections. (Scale bar 50 µm)

**Table 1.**
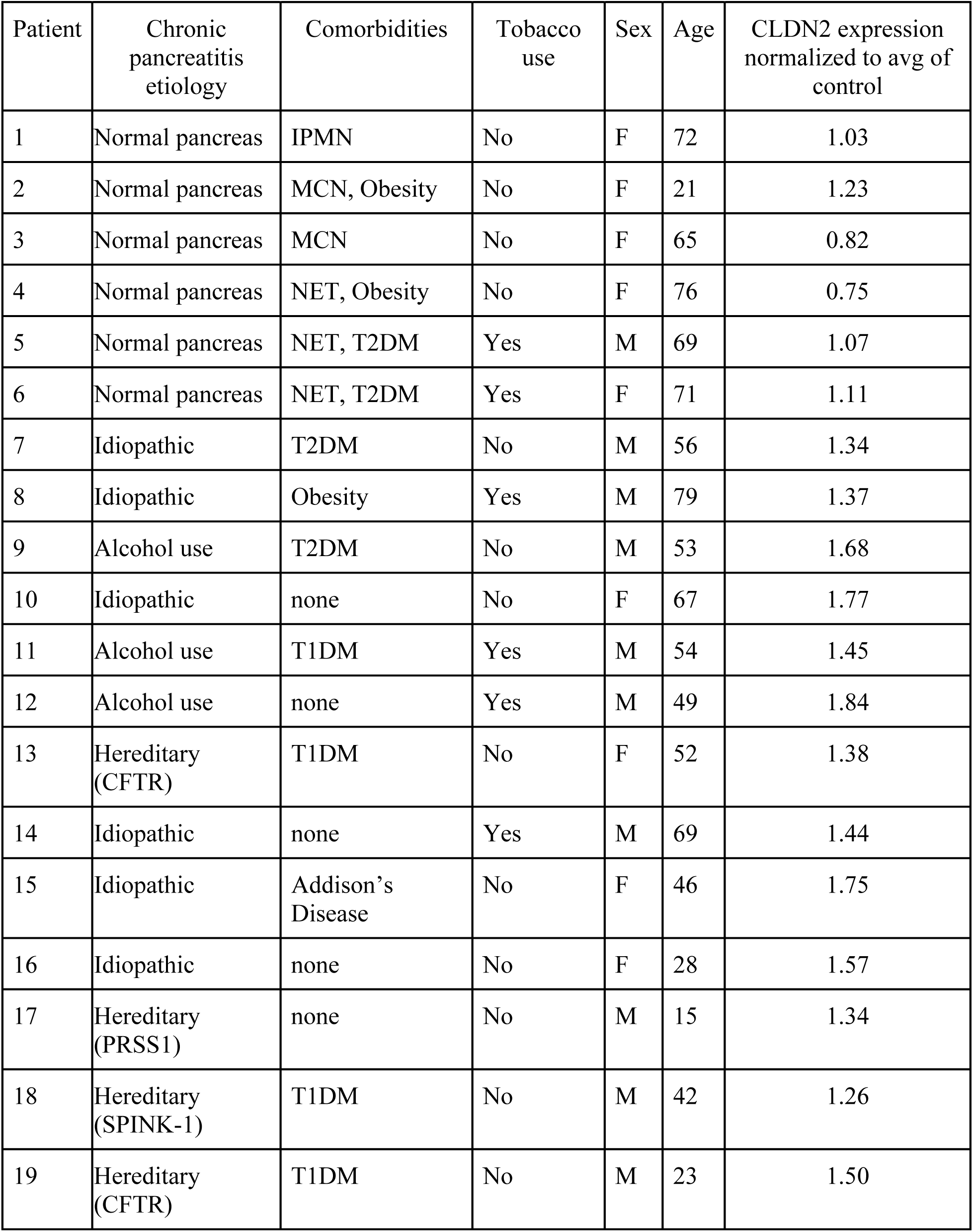
Patient characteristics. **Human chronic pancreatitis health history and CLDN2 quantification.** Patients 1-6: Normal pancreas sampled at margins or far from benign pancreatic neoplasms. (IPMN: intraductal pancreatic mucinous neoplasm, MCN: mucinous cystic neoplasm, NET: well differentiated pancreatic neuroendocrine tumor) Patients 7-19: Patients with chronic pancreatitis not associated with neoplasms.

| Patient | Chronic pancreatitis etiology | Comorbidities | Tobacco use | Sex | Age | CLDN2 expression normalized to avg of control |
| --- | --- | --- | --- | --- | --- | --- |
| 1 | Normal pancreas | IPMN | No | F | 72 | 1.03 |
| 2 | Normal pancreas | MCN, Obesity | No | F | 21 | 1.23 |
| 3 | Normal pancreas | MCN | No | F | 65 | 0.82 |
| 4 | Normal pancreas | NET, Obesity | No | F | 76 | 0.75 |
| 5 | Normal pancreas | NET, T2DM | Yes | M | 69 | 1.07 |
| 6 | Normal pancreas | NET, T2DM | Yes | F | 71 | 1.11 |
| 7 | Idiopathic | T2DM | No | M | 56 | 1.34 |
| 8 | Idiopathic | Obesity | Yes | M | 79 | 1.37 |
| 9 | Alcohol use | T2DM | No | M | 53 | 1.68 |
| 10 | Idiopathic | none | No | F | 67 | 1.77 |
| 11 | Alcohol use | T1DM | Yes | M | 54 | 1.45 |
| 12 | Alcohol use | none | Yes | M | 49 | 1.84 |
| 13 | Hereditary (CFTR) | T1DM | No | F | 52 | 1.38 |
| 14 | Idiopathic | none | Yes | M | 69 | 1.44 |
| 15 | Idiopathic | Addison's Disease | No | F | 46 | 1.75 |
| 16 | Idiopathic | none | No | F | 28 | 1.57 |
| 17 | Hereditary (PRSS1) | none | No | M | 15 | 1.34 |
| 18 | Hereditary (SPINK-1) | T1DM | No | M | 42 | 1.26 |
| 19 | Hereditary (CFTR) | T1DM | No | M | 23 | 1.50 |

### CLDN2 expression is increased in caerulein-induced experimental pancreatitis in mice

We next sought to understand how upregulated CLDN2 expression may impact pancreatitis severity by using mouse models. We first determined if the modulation of CLDN2 expression can be recapitulated in these models. Experimental pancreatitis was induced by repeated injection of the cholecystokinin homolog caerulein (first injection at day 0). Following eight hourly caerulein injections per day for two days, pancreatic samples were collected 1d after the final injection (day 2), representing the early phase of caerulein-induced experimental pancreatitis and 7 d after the final injection (day 8), representing the late phase of the disease (Fig. 2A). H&E staining showed that tissues obtained during early and late phases of disease exhibited histological evidence of tissue damage, including mild expansion of the interstitial space due to edema and inflammation (Fig. 2B). In the late phase of disease, inflammation and fibrin deposition was more predominant, particularly in periductal areas (Fig. 2B). Immunohistochemical staining showed CLDN2 expression was increased in the ductal epithelial cells in both the early (p=0.0145), and late (p=0.0049) phases of disease (Fig. 2C-D). CLDN2 was increased at the tight junction, cell membranes, and in cytoplasmic vesicles. These findings demonstrate that CLDN2 is consistently upregulated in both human and experimental pancreatitis, reinforcing its potential contribution to disease development and highlighting the value of the caerulein-induced mouse model for further mechanistic studies.

**Figure 2.**
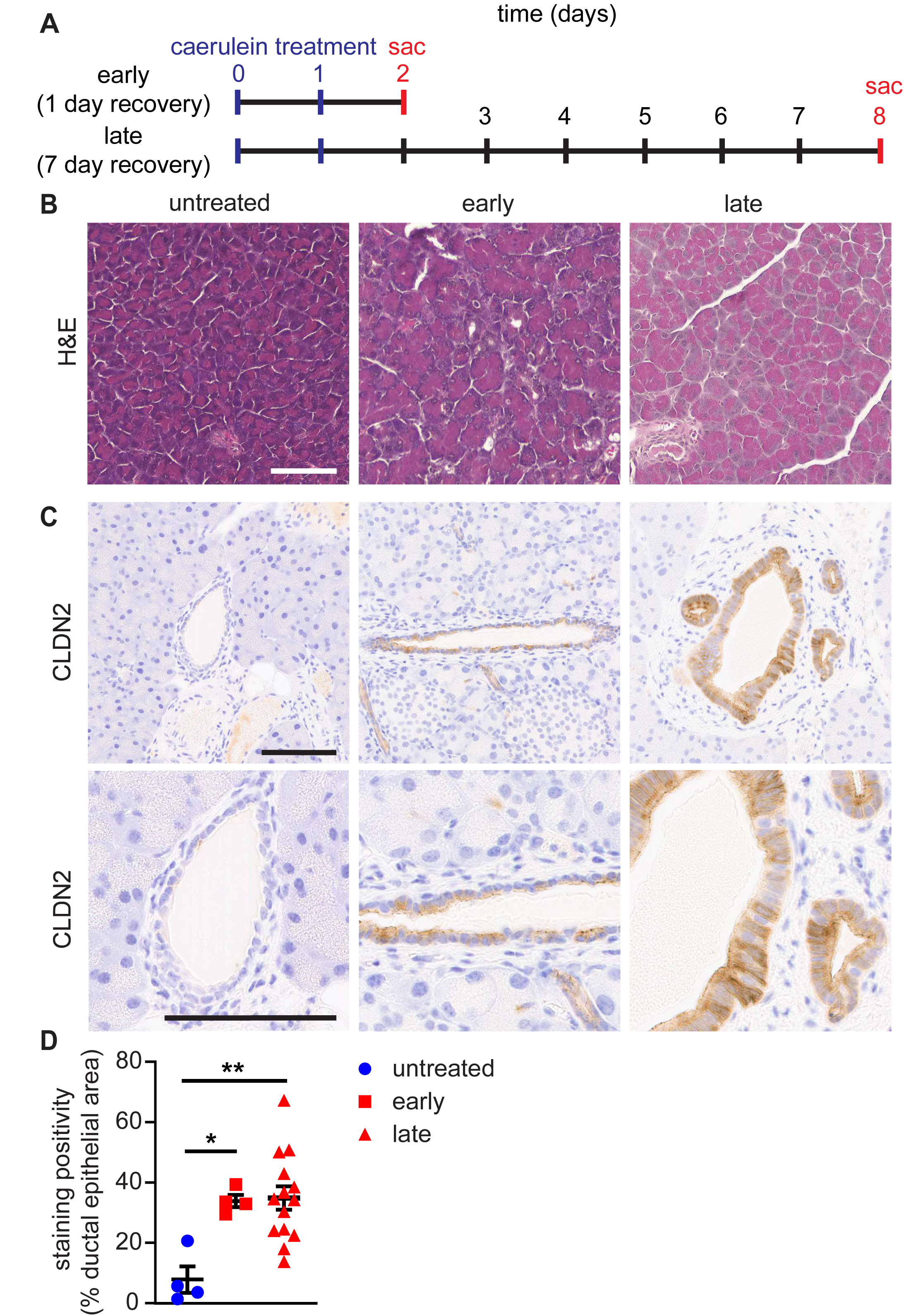
CLDN2 protein expression is increased in caerulein-induced experimental pancreatitis. **A.** Timeline of caerulein-induced experimental pancreatitis. Following 8 hourly caerulein injections daily for two days, pancreata were harvested during early phase (1 d recovery) and late phase (7 d recovery) of the disease. **B.** Representative images of H&E-stained mouse pancreatic tissue sections. (Scale bar 100 µm) **C.** CLDN2 staining of mouse pancreatic tissue sections. (Scale bar 100 µm) **D.** Quantification of CLDN2 staining as percent positivity (positive pixels/total pixels in pancreatic ductal epithelium) in pancreatic ducts. Blue circles: untreated mouse pancreas tissue (n=4), red squares: 1 d recovery following caerulein injection (n=4), red triangles: 7 d recovery following caerulein injection (n=8). Two-tailed one-way ANOVA with Bonferroni *post hoc* analysis was used for statistical analysis. *p<0.05 and **p<0.01. Untreated versus 1d recovery mice: p=0.015, untreated versus 7 d recovery: p=0.0049.

### IFNγ drives claudin-2 expression in pancreatic ductal epithelial cells

Our data showed CLDN2 is upregulated in pancreatic ductal epithelial cells during chronic pancreatitis. We hypothesized that CLDN2 upregulation could be influenced by various cytokines, including IL-13, IL-6, IL-22, TNFα, and IFNγ, which are known to increase its expression in other tissues (Prasad et al., 2005; Raju et al., 2020; Suzuki et al., 2011; Tsai et al., 2017b; Weber et al., 2010; Zhang et al., 2017). To evaluate the effect of these cytokines on CLDN2 expression in pancreatic ductal epithelium, we treated pancreatic ductal epithelial organoids derived from C57BL/6 mice with cytokines. In contrast to previous pancreatic ductal adenocarcinoma cell line derived 2D culture models, our non-transformed 3D pancreatic ductal epithelial organoid model provides a system that better reflects normal pancreatic ductal physiology. Out of these cytokines, only IFNγ increased *Cldn2* mRNA expression (Fig. 3A). Further study showed that IFNγ increases *Cldn2* mRNA expression in a dose-dependent manner (Fig. 3B), as determined by qRT-PCR. CLDN2 protein expression was also increased in organoids treated with IFNγ, demonstrated by IHC staining of organoid sections (Fig. 3C). To determine if IFNγ can induce CLDN2 expression *in vivo*, we injected IFNγ intraperitoneally and collected mouse pancreata 24 hour later. qRT-PCR and IHC staining showed that IFNγ significantly induced CLDN2 expression at both mRNA and protein level (Fig. 3D-E). Together, these data show that IFNγ can induce CLDN2 expression both *in vitro* and *in vivo*. To further explore this role, we evaluated whether CLDN2 upregulation in the caerulein-induced mouse model of pancreatitis can be impacted in IFNγ knockout (*Ifng* KO) mice. While CLDN2 expression still increased in *Ifng* KO mice at day 2 following initiation of caerulein treatment, this increase was significantly lower than in wildtype mice treated with caerulein (Fig. 3F). Collectively, these data support a role for IFNγ in driving CLDN2 expression in pancreatic ductal epithelial cells.

**Figure 3.**
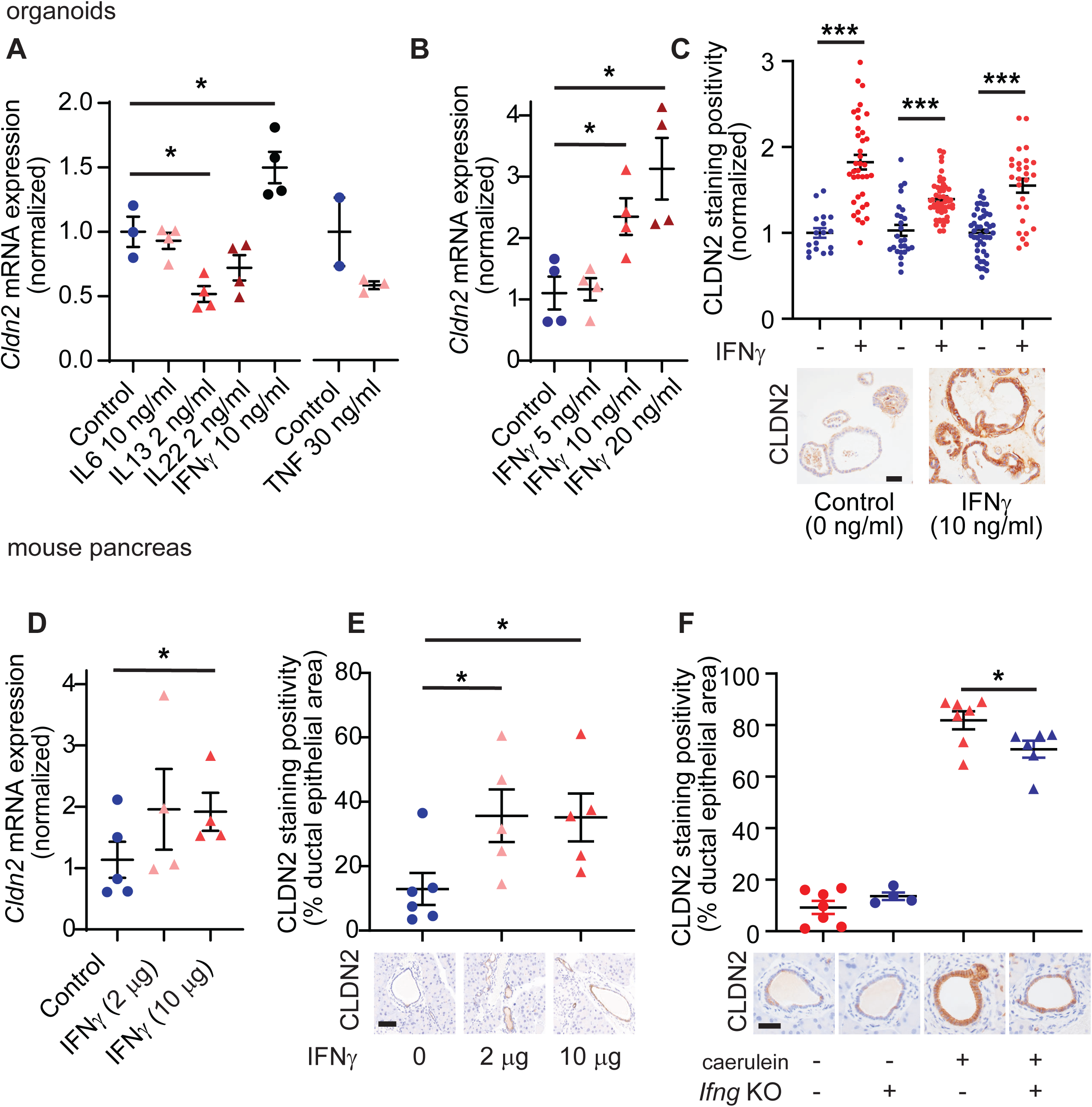
IFNγ upregulates claudin-2 mRNA and protein expression *in vitro* and *in vivo*. **(A)** *Cldn2* mRNA expression was measured in pancreatic organoids treated with various cytokines—IL-6 (10 ng/mL), IL-13 (2 ng/mL), IL-22 (2 ng/mL), and IFNγ (10 ng/mL)—for 48 hours to assess their differential effects on *Cldn2* induction. Statistical analysis was performed using a two-tailed one-way ANOVA with Bonferroni correction for post hoc comparisons, with a significance threshold of *p*<0.05. In a separate experiment, TNF treatment (30 ng/mL for 24 hours) showed no effect on *Cldn2* expression, as determined by Student’s t-test. **(B)** *Cldn2* mRNA expression in pancreatic organoids treated with varying doses of IFNγ (5, 10, and 20 ng/mL), showing dose-dependent upregulation. Two-tailed one-way ANOVA with Bonferroni corrected *post hoc* analysis was used for statistical analysis. *p<0.05 **(C)** IFNγ-treated organoids display increased CLDN2 protein expression, as indicated by increased staining in ductal epithelial cells. Unpaired Student’s *t* test was performed on three separate experiments, ***p<0.001. scale bar 50 µm **(D)** *Cldn2* mRNA expression in mouse pancreas tissue following interperitoneal IFNγ injection (2 µg or 10 µg). Two-tailed one-way ANOVA with Bonferroni corrected post hoc analysis was used for statistical analysis. *p<0.05. **(E)** IHC images of CLDN2 showing increased protein expression in mouse pancreas tissue following intraperitoneal IFNγ injection (2 µg or 10 µg). Two-tailed one-way ANOVA with Bonferroni post hoc analysis was used for statistical analysis. scale bar 100 µm **(F)** IHC images of CLDN2 showing caerulein-pancreatitis-induced upregulation of claudin-2 in pancreatic ducts (1 d recovery) is diminished in Ifng knockout mice. Two-tailed one-way ANOVA with post hoc analysis was used. *p<0.05. scale bar 50 µm.

### Cldn2 KO exacerbates tissue inflammation in the early phase of caerulein-induced experimental pancreatitis

Although studies above demonstrated CLDN2 is upregulated during pancreatitis development, the functional consequences for such upregulation remain unknown. To determine the impact of CLDN2 on pancreatitis development, we assessed how the loss of CLDN2 expression affects the progression of caerulein-induced pancreatitis by using *Cldn2* KO mice. *Cldn2* KO mice exhibit calcium deposits in the kidney after 4.5 months, but do not exhibit features of systemic disease or weight change in an unstressed state and their pancreata appear histologically normal by H&E stain (Curry et al., 2020; Muto et al., 2010).

Both control and *Cldn2* KO mice had similar weight change overtime (Supplementary Fig. 2A). In the early phase (1 d after last caerulein injection) of pancreatitis, H&E staining showed more significant tissue reorganization in *Cldn2* KO mice following caerulein treatment (Fig. 4A), and immunohistochemical staining showed that while there was very low CLDN2 staining in pancreas tissues in control and none in *Cldn2* KO mice without treatment, caerulein only induced significant CLDN2 staining increase in control mice (Fig 4B). In this phase, little tissue fibrosis occurred in either control or *Cldn2* KO mice, as indicated by limited blue staining in the trichrome stained tissue (Fig. 4C, E). Immunohistochemical staining of CD45, a pan-leukocyte marker, was performed to assess tissue inflammation. While the percent of total cells staining positive for CD45 in pancreata from both control and *Cldn2* KO mice increased following caerulein treatment, the number increased significantly more in *Cldn2* KO mice (Fig. 4D, F, n=8 and n=8 for control and *Cldn2* KO groups, respectively). This indicates CLDN2 limits tissue inflammation in the early phase of caerulein-induced experimental pancreatitis.

**Figure 4.**
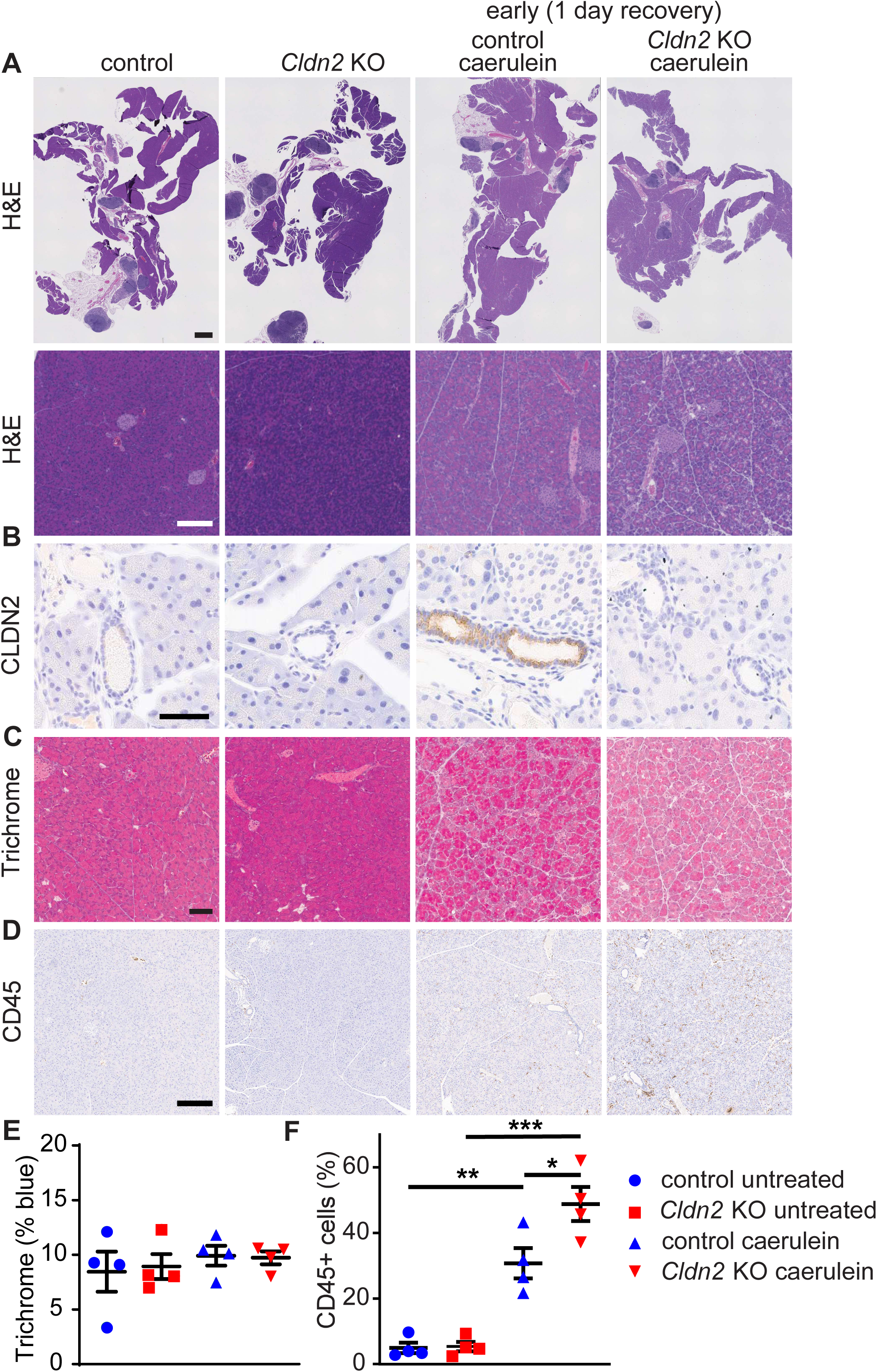
*Cldn2* KO mice have increased tissue leukocytes in early phase of experimental pancreatitis. Pancreatic tissues were harvested 1 d following caerulein injection. **A.** Representative H&E-stained mouse pancreatic tissue sections. (Top panels: scale bar 1 mm, bottom panels: scale bar 200 µm) **B.** CLDN2 staining of mouse pancreas tissue sections. (Scale bar 50 µm) **C.** Trichrome staining of mouse pancreatic tissue sections. (Scale bar 100 µm) **D.** CD45 staining of leukocytes in mouse pancreatic tissue sections. (Scale bar 200 µm) **E.** Quantification of percent collagen/fibrosis staining (blue) area over total tissue area, visualized by trichrome staining. Blue circles: control mice (n=4), red squares: *Cldn2* KO mice (n=4), blue triangles: caerulein treated control mice (n=4), red upside down triangles: caerulein treated *Cldn2* KO mice (n=4) (all comparisons not significant p>0.05). **F.** Quantification of CD45 positive cells. Blue circles: control mice (n=4), red squares: *Cldn2* KO mice (n=4), blue triangles: caerulein treated control mice (n=4), red upside down triangles: caerulein treated *Cldn2* KO mice (n=4) (control untreated vs. control caerulein p=0.0012, *Cldn2* KO untreated vs. *Cldn2* KO caerulein p=0.000009; control caerulein vs. *Cldn2* KO caerulein p=0.017). Two-tailed one-way ANOVA with Bonferroni *post hoc* analysis was used for statistical analysis. **p<0.01, and ***p<0.001.

### Cldn2 KO exacerbates pancreatic fibrosis in late phase of caerulein-induced experimental pancreatitis

We next evaluated how the lack of CLDN2 expression affects the late phase (7 d after last caerulein injection) of experimental pancreatitis. At this phase, there was no significant difference in weight between the groups (Supplementary Fig. 2B). In contrast, there was still significant tissue damage differences between control and *Cldn2* KO mice (Fig. 5A). The pancreata from the *Cldn2* KO mice showed greater cellular vacuolization, edema, inflammation, and acinar cell dedifferentiation. Trichrome staining showed that despite limited fibrosis in control mice, there was significant tissue fibrosis in *Cldn2* KO mice (n=14 and 17 for control and *Cldn2* KO groups, respectively; p=0.0072) (Fig. 5A-C, Supplementary Fig. 3). CD45 immunohistochemical staining showed the percent of CD45 positive leukocytes in this later phase was lower in both groups of mice, with no statistical significance in both groups, although there was still a small number of *Cldn2* KO mice that had high numbers of tissue leukocytes. Thus, CLDN2 limits tissue damage and fibrosis in the late phase of experimental pancreatitis.

**Figure 5.**
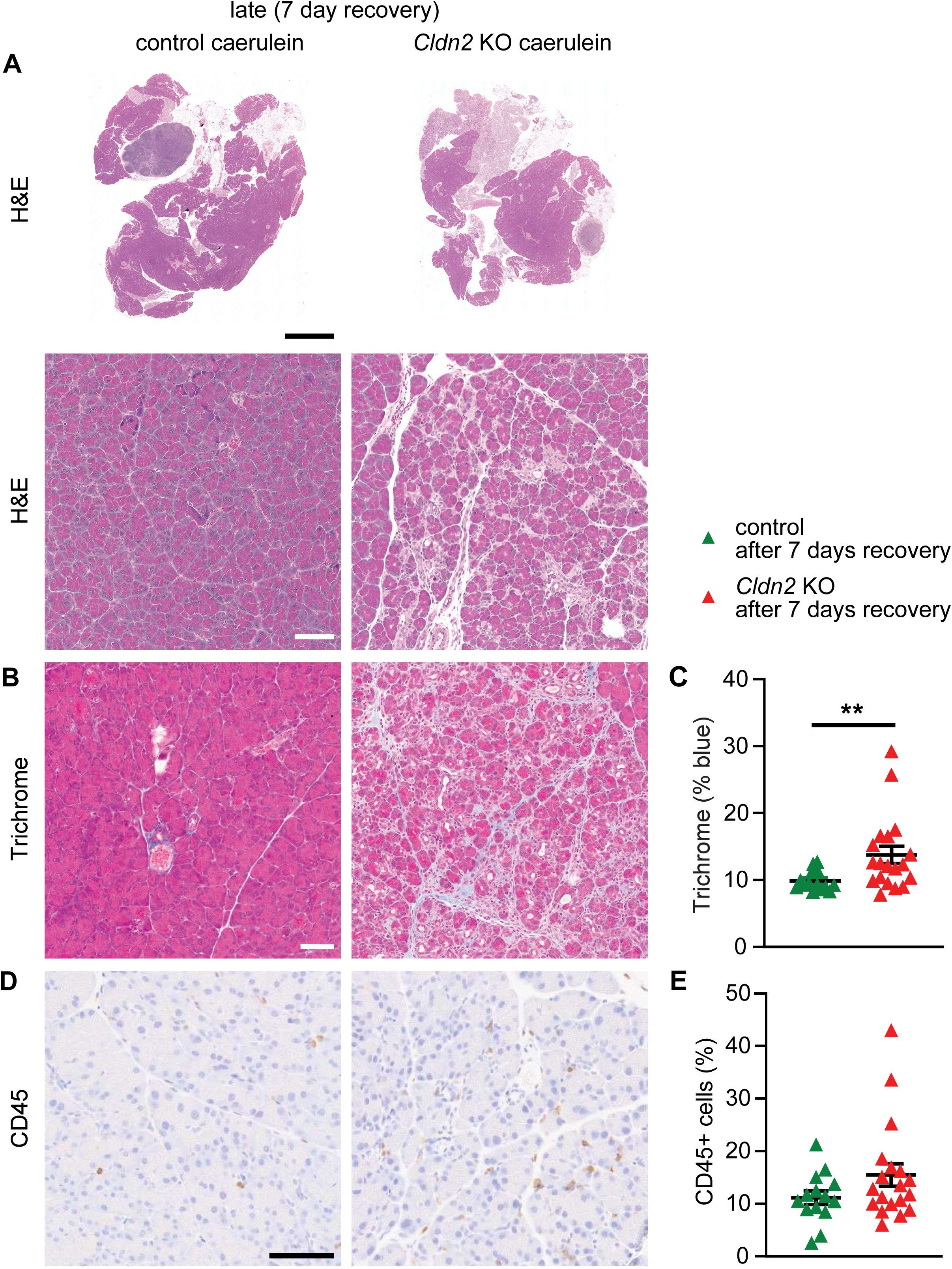
*Cldn2* KO mice have increased tissue fibrosis during the late phase of experimental pancreatitis. **A.** Representative H&E-stained mouse pancreatic tissue sections. (Top panels: scale bar 2 mm, bottom panels: scale bar 100 µm) **B.** Trichrome staining of mouse pancreatic tissue sections. (Scale bar 100 µm) **C.** Quantification of percent collagen/fibrosis staining (blue) area over total tissue area, visualized by trichrome staining. Green triangles: caerulein injected control mice (n=14), red triangles: caerulein injected *Cldn2* KO mice (n=17) (p=0.0072)**. D.** CD45 staining of leukocytes in mouse pancreatic tissue sections. (Scale bar 100 µm) **E**. Quantification of CD45positive cells. Green triangles: caerulein injected control mice (n=14), red triangles: caerulein injected *Cldn2* KO mice (n=17) (p=0.0942). Two-tailed unpaired Student’s *t* test was used for statistical analysis. **p<0.01.

### CLDN2 regulates pancreatic ductal epithelial transport

CLDN2 is primarily expressed in pancreatic ductal epithelial cells, and its expression is upregulated in this cell type in the setting of pancreatitis. Furthermore, CLDN2 can form a paracellular channel that is selective for small cations and can also mediate water transport, based on studies on intestinal and renal epithelium (Muto et al., 2010; Tsai et al., 2017b). Thus, it is possible that CLDN2 can also regulate small cation and water transport in pancreatic ductal epithelium. To test this possibility, we generated pancreatic ductal epithelial organoids from control and *Cldn2* KO mice. Immunofluorescent staining of formalin fixed organoid sections confirmed that CLDN2 is expressed in control organoids, while no CLDN2 expression can be observed in *Cldn2* KO organoids (Fig. 6A).

**Figure 6.**
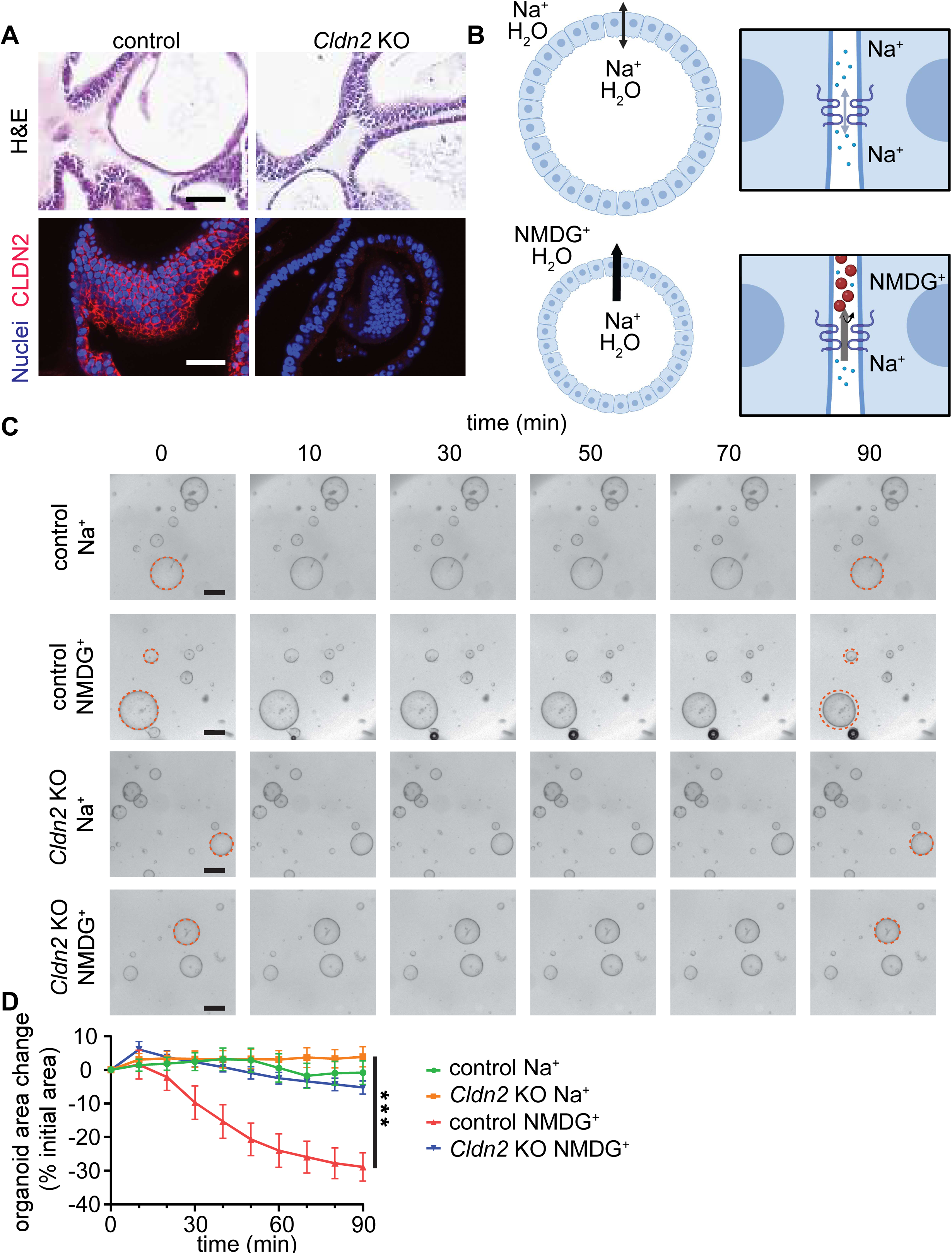
CLDN2 mediates Na^+^-gradient-driven water transport in pancreatic ductal epithelium. **A.** Representative H&E and CLDN2 staining of control and *Cldn2* KO pancreatic ductal epithelial organoid sections. (Scale bars 50 µm) **B.** Schematic presentation of possible mechanism of Na^+^ gradient-driven organoid shrinkage. Top panels: no unidirectional water transport when luminal and extraluminal solution both contain Na^+^. Bottom panels: when extraluminal Na^+^ is substituted with NMDG^+^, a monovalent cation driving force exists for water to move out of organoid lumen. **C.** Representative time lapse images of organoid within the same field over time for each treatment condition. Dashed red circles indicate initial organoid outline. Substitution of Na^+^ in extraluminal imaging media (HBSS) with isomolar Na^+^ (first and third rows) or NMDG^+^ (second and fourth rows) occurred at t=0. (Scale bars 50 µm) **D.** Quantification of normalized organoid area. Green circles and line: control organoids mock changed to Na^+^ containing HBSS (n=5), orange squares and line: *Cldn2* KO organoids mock changed to Na^+^ containing HBSS (n=9), red triangles and line: control organoids changed to NMDG^+^ containing HBSS (n=5), upside down blue triangles and line: *Cldn2* KO organoids changed to NMDG^+^ containing HBSS (n=10) (p<0.0001 among different groups). Repeated measures 2-way ANOVA test. ***p<0.001.

We hypothesized that if a sodium selective pathway is important for pancreatic ductal epithelial transport, creating an iso-osmotic Na^+^ gradient across the epithelial layer of a pancreatic organoid would create a driving force for unidirectional Na^+^ transport. If Na^+^ flows through such a selectively permeable pathway, water must follow to maintain ionic equilibrium (Fig. 6B). To test this, we substituted Na^+^ in extraluminal imaging media with isomolar CLDN2 impermeable large organic ion N-methyl-D-glutamine (NMDG^+^), creating a high intraluminal to extraluminal Na^+^ concentration gradient (Fig. 6B). While changing regular imaging media with the same Na^+^ containing media had little effect on organoid size, substitution with iso-osmotic NMDG^+^ containing Na^+^ free media caused progressive decrease in organoid size (Fig. 6C, D row 2 vs. row 1). In contrast, when such ion substitution was performed on *Cldn2* KO organoids, such size change was limited (Fig. 6C, D row four vs. row 3). This shows that a Na^+^ driven fluid transport pathway exists in pancreatic ductal epithelium, which is CLDN2 dependent.

### CLDN2 participates in secretagogue-induced pancreatic ductal epithelial secretion

Having identified that CLDN2 can regulate Na^+^-dependent transport in pancreatic ductal epithelium, we next sought to assess contributions of CLDN2 to active pancreatic ductal secretion using a forskolin induced organoid swelling assay *(Dekkers et al., 2013; Molnár et al*., 2020). In this assay, forskolin activates adenylate cyclase and induces cystic fibrosis transmembrane regulator (CFTR)-mediated cellular secretion of Cl^-^ into the organoid lumen (Fig. 7A). This generates an electrochemical gradient that drives Na^+^ entry into the lumen *(Madácsy et al*., 2018). In control pancreatic ductal epithelial organoids, forskolin induced an increase in organoid size. (Fig. 7B, C). In contrast, forskolin only caused a limited increase in *Cldn2* KO organoid size. These data further show a functional paracellular Na^+^ permeable CLDN2 channel is required for pancreatic ductal epithelial secretion. IHC staining showed CFTR expression was maintained in *Cldn2* knockout organoids and tissues, indicating the defect in forskolin-induced swelling in *Cldn2* KO organoids is not caused by decreased CFTR expression (Supplementary Fig. 4A and 4B).

**Figure 7.**
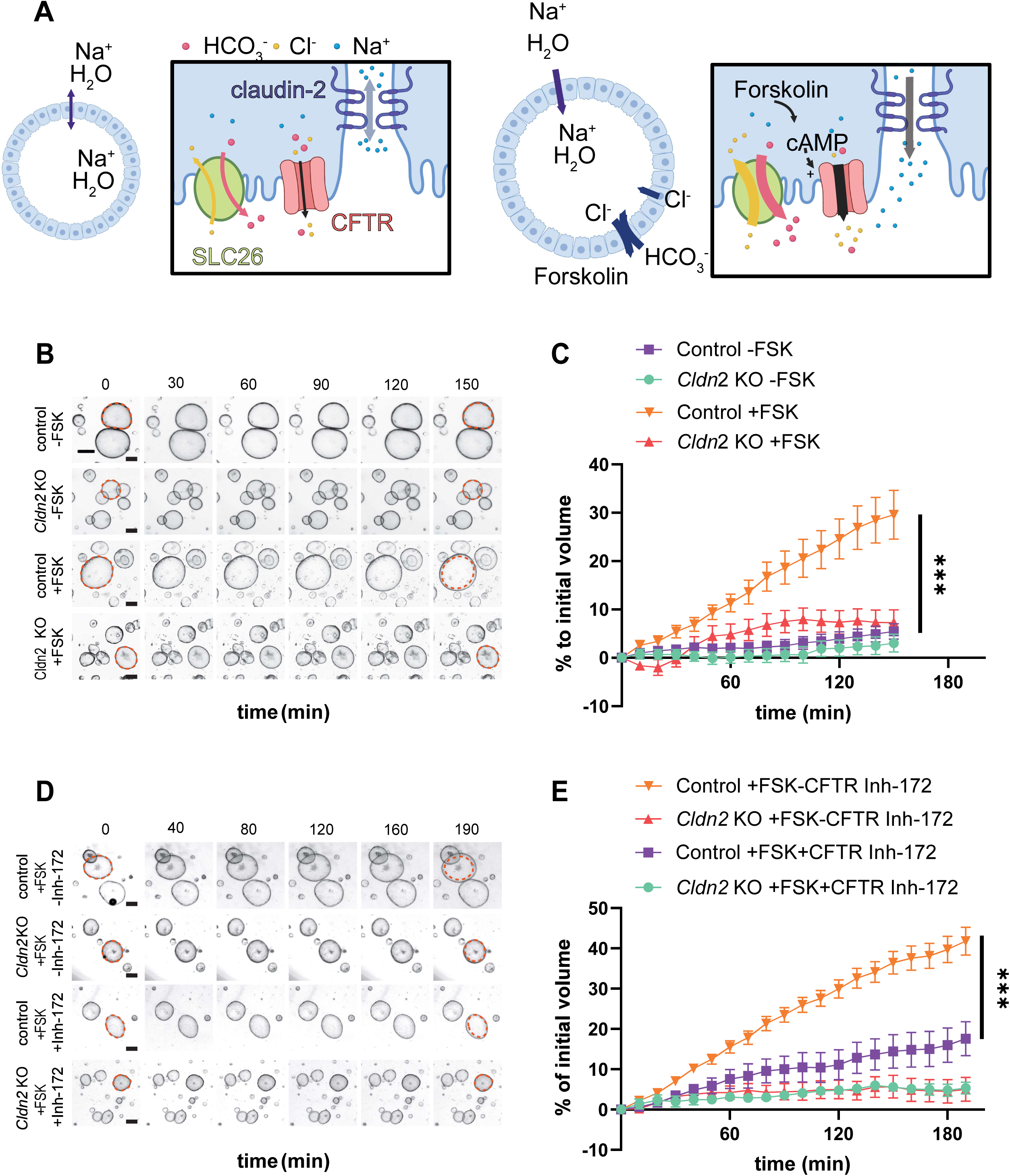
CLDN2 mediates forskolin-induced mouse pancreatic ductal organoid swelling. **A.** Schematic presentation of possible mechanism of forskolin (FSK) induced organoid swelling. Left panels: no unidirectional water transport occurs at resting state. Right panels: forskolin treatment activates CFTR to induce Cl^-^ secretion. This leads to SLC26 mediated Cl^-^/HCO3^-^ exchange, resulting in HCO3^-^ secretion. This drives paracellular Na^+^secretion. This creates an osmotic gradient for water secretion. **B.** Representative time lapse images of control and *Cldn2* KO mouse organoids with and without forskolin (FSK) treatment. Red circles indicate initial organoid outline. (Scale bar 50 µm) **C.** Quantification of normalized organoid area with and without FSK stimulation. Green squares: control organoids without FSK treatment (n=11), blue circles: *Cldn2* KO organoids without FSK treatment (n=24), orange triangles: control organoids with 10 < M FSK treatment (n=12), red triangles: *Cldn2* KO organoids with 10 < M FSK treatment (n=9) (p<0.0001). Repeated measures 2-way ANOVA test with. ***p<0.001. **D**. Representative time lapse images of control and *Cldn2* KO mouse organoids with FSK and with and without CFTR Inhibitor-172 treatment. Red circles indicate initial organoid outline. (Scale bar 50 µm) **E**. Quantification of normalized organoid area for forskolin-induced swelling of control and Cldn2 KO organoids in the presence or absence of CFTR Inhibitor-172. Orange triangles: control organoids treated with FSK without CFTR inhibitor-172 (n=20); red triangles: *Cldn2* KO organoids treated with FSK without CFTR inhibitor (n=7); purple squares: control organoids treated with FSK and CFTR Inhibitor-172 (n=9); green circles: *Cldn2* KO organoids treated with FSK and CFTR Inhibitor-172 (n=15). Repeated measures 2-way ANOVA test with. ***p<0.001.

To further determine the contribution of CFTR to forskolin-induced organoid swelling, we employed the CFTR inhibitor, CFTR inh-172. Inhibiting CFTR significantly reduced forskolin-induced swelling in WT pancreatic ductal organoids, demonstrating forskolin-induced organoid swelling depends on CFTR-mediated Cl^-^ secretion. In contrast, forskolin-induced swelling remained negligible in the *Cldn2* knockout organoids, and CFTR inh-172 treatment of *Cldn2* knockout organoids did not result in any significant difference in organoid size change compared to forskolin treatment alone.

Taken together, these data underscore a critical role of CLDN2 facilitated paracellular Na^+^ permeability for pancreatic ductal epithelial secretion. These studies also show CFTR activity alone is insufficient to induce significant organoid swelling in the absence of CLDN2, highlighting that an interplay between CLDN2 and CFTR is required for fluid secretion in pancreatic ducts.

## DISCUSSION

Pancreatitis is an inflammatory disease of the pancreas that originates from a complex interplay of genetic predispositions, environmental triggers, autoimmune diseases, and metabolic disorders (Beyer *et al*., 2020). Aberrant activation of digestive enzymes within the pancreas is thought to play a major role in the development of pancreatitis. This is supported by evidence of genetic mutations implicated in chronic pancreatitis, which often affect the trypsin-dependent pathway. Genes such as *PRSS1*, *SPINK1*, and *CTRC* are known to contribute to pancreatitis development through these mutations. Genes that regulate ion and fluid balance in the pancreatic ducts, such as *CFTR, CASR,* and *TRPV6* also impact pancreatitis development (Masamune et al., 2020; Mounzer & Whitcomb, 2013). Defects in these ductal associated genes may limit pancreatic fluid secretion, increasing digestive enzyme concentration, leading to autodigestion of the pancreas. This suggest that other proteins that regulate pancreatic fluid secretion may also impact pancreatitis development. Consistent with this notion, GWAS analyses identified a gene that contributes to paracellular cation transport and epithelial barrier function, CLDN2, as a risk gene for pancreatitis (Deng & Li, 2020; Giri et al., 2016; Weiss et al., 2018; Whitcomb et al., 2012).

CLDN2 belongs to the claudin family of the tetraspanning transmembrane tight junction proteins. It is well established as a cation selective ion channel and has also been implicated to facilitate water transport in the same direction of ion gradient, in a “solvent drag” fashion. In the intestines, CLDN2 is thought to participate in intestinal crypt secretion. Interestingly, while *Cldn2* knockout (KO) enhances susceptibility to intestinal bacterial infection, it appears to limit experimental inflammatory bowel disease (Raju et al., 2020; Tsai et al., 2017a).

Thus, the same CLDN2 governed physiological process may differentially regulate distinct disease processes in a context dependent manner. In the case of pancreatitis, from a pathophysiological perspective, it is conceivable CLDN2 alterations could affect pancreatitis development. However, it is not clear if upregulated or decreased CLDN2 function promotes pancreatitis development. Results of GWAS studies do not readily provide an answer, as all *CLDN2* associated SNPs exist outside of protein coding regions of *CLDN2*. Thus, they do not affect CLDN2 amino acid sequence but are more likely to alter its expression. Although an initial publication connected a pancreatitis associated SNP (*rs12688220*) within the *CLDN2*-*MORC4* region to altered CLDN2 protein localization within the pancreatic ductal epithelial cells, subsequent analysis suggested this locus is associated with *MORC4*, rather than *CLDN2*. Thus, it remains unclear if increased or decreased CLDN2 function may promote pancreatitis development. To address this question, we first determined if CLDN2 expression is altered in pancreatitis.

Using pancreatic specimens from patients with chronic pancreatitis of various etiologies, we demonstrated that CLDN2 expression is upregulated, particularly in pancreatic ductal cells. CLDN2 expression is also elevated in both the early and late phases of caerulein-induced pancreatitis in mice, indicating that this may be a useful model for determining CLDN2’s contribution to pancreatitis development. In this model, *Cldn2* deletion exacerbated disease development, with increased immune cell infiltration and tissue fibrosis. Thus, upregulated CLDN2 expression in pancreatitis, is a compensatory mechanism that limits pancreatitis development. This also points to the possibility that enhancing CLDN2 function may serve as a novel method for pancreatitis treatment.

With these core findings, it is necessary to understand how CLDN2 may be upregulated in pancreatitis. It is known that CLDN2 upregulation is associated with tissue inflammation across various conditions. For example, in the gastrointestinal tract, CLDN2 expression is increased in a broad range of diseases, including inflammatory bowel disease, (Heller et al., 2005; Prasad et al., 2005; Weber et al., 2010) infectious colitis, (Liu et al., 2013) and celiac disease (Szakál et al., 2010). In the lungs, a similar pattern of CLDN2 upregulation has been observed under chronic inflammatory conditions (Kaarteenaho-Wiik & Soini, 2009; Zou et al., 2020). These findings suggest that CLDN2 upregulation is a common response to chronic inflammation in various organs. In the gastrointestinal tract, CLDN2 upregulation has been reported to be driven by many different inflammatory cytokines (Amoozadeh *et al*., 2015; Mankertz *et al*., 2009; Suzuki *et al*., 2011; Tian *et al*., 2018; Wang *et al*., 2017; Weber *et al*., 2010; Yamamoto *et al*., 2004). Using normal pancreatic ductal epithelial organoid model, our study showed that IFNγ is an effective inducer of CLDN2 expression at the transcriptional level, while the other cytokines tested, including IL-13, IL-6, IL-22, and TNFα, do not increase *Cldn2* mRNA expression. When injected to wild type mice, IFNγ also induced CLDN2 expression at both the mRNA and protein levels.

These findings may be functionally significant, as *Ifnγ* KO mice have been reported to have more severe caerulein-induced experimental pancreatitis (Hayashi et al., 2007). In our study, caerulein-induced CLDN2 expression is reduced in *Ifng* KO mice. Although our data indicate that IFNγ is not the only driver to upregulate CLDN2 expression, they suggest that IFNγ limits pancreatitis development at least partially through inducing CLDN2 expression.

To completely understand how CLDN2 is upregulated to limit pancreatitis development, it is necessary to understand how other factors may drive CLDN2. It is unlikely that IL-13, IL-6 and TNF are major drivers, not only because our studies using organoids demonstrate they do not induce *Cldn2* mRNA expression, but previous studies using genetically modified animals demonstrate these cytokines promote, rather than limit, experimental pancreatitis development (Denham et al., 1997; Suzuki et al., 2011; Xue et al., 2015). Additional studies are needed to systemically define regulators for CLDN2 expression in the context of pancreatitis development. We further sought to understand how CLDN2 expression may protect pancreatitis development. Because CLDN2 can regulate epithelial tight junction cation and water transport, we determined the ability of CLDN2 to impact fluid transport in pancreatic ducts by using normal pancreatic duct epithelial organoid derived from control and *Cldn2* KO mice (Molnár et al., 2020; Varga et al., 2023). This model is superior to 2D cultured pancreatic ductal adenocarcinoma-derived transformed ductal epithelial, as it maintains near physiological general organization of ductal epithelial cells, proper cell polarity, and expression of epithelial transport machinery, without mutations found in cancer cells.

Additionally, this model system provides greater throughput and experimental flexibility compared to isolated pancreatic ducts. Our organoid-based experiments provided novel insights into pancreatic ductal epithelial transport. Substituting extraluminal Na^+^ with NMDG^+^ caused a progressive decrease in organoid size, indicating net fluid extrusion from the lumen. Since water movement in our system depends on local osmotic gradients, these observations suggest an asymmetry in Na^+^ and NMDG^+^ transport across the epithelium. This indicates a size-selective cation-permeable pathway in the pancreatic ductal epithelium that drives fluid transport. Because this pathway was largely absent in *Cldn2* KO organoids, and CLDN2 mediates selective cation transport, our data suggest that a Na+ gradient primarily drives fluid transport through a CLDN2-dependent paracellular pathway.

During pancreatic secretion, CFTR mediated Cl^-^ secretion, and the subsequent Cl ^-^ /HCO3^-^ exchange through SLC26 family members drives HCO3^-^ and fluid secretion in the pancreatic ducts (Pallagi *et al*., 2015). It is well recognized that a paracellular shunt pathway is required for Na^+^ to follow anion secretion (Frizzell & Schultz, 1972). However, the molecular entity that creates this paracellular passive Na ^+^ permeable pathway is unknown. In addition to the result discussed in the above paragraph demonstrating the presence of a CLDN2 dependent Na^+^ permeable pathway, our forskolin-induced swelling assay further suggests paracellular Na^+^ transport during pancreatic ductal secretion is also mediated by CLDN2. Our studies using a CFTR inhibitor directly show that CFTR activity is required for CLDN2 dependent swelling of pancreatic ductal epithelial organoids, reinforcing the notion that both transcellular (CFTR) and paracellular (CLDN2) transport are both required for pancreatic ductal secretion.

Although CLDN2 has also been reported to function as a water transporter, and our data show loss of CLDN2 function can limit fluid transport in both organoid models used in this study, it is not clear if observed water transport is directly mediated by CLDN2. It is possible that CLDN2 controlled directional paracellular Na^+^ transport only provides an osmotic gradient that drives water transport through other channels, including aquaporins. It is also possible that both passive Na^+^ and water transport are directly mediated by CLDN2. Relative contributions of water transport directly through CLDN2 channels and other water transporters remain to be investigated in the future.

Our studies using chronic pancreatitis patient samples and caerulein-induced pancreatitis in mice showed that CLDN2 expression is upregulated in pancreatic ductal epithelium during inflammation. Given results obtained from organoid based studies, such CLDN2 upregulation can facilitate ion and water secretion into the lumen during pancreatitis, and this could be a compensatory pathophysiological change to limit pancreatitis. In the caerulein-induced pancreatitis model, the absence of this upregulation, such as in *Cldn2* KO mice, results in paracellular Na^+^ secretion into the ductal lumen being reduced. This would create a higher luminal Cl^-^ concentration, limiting further Cl^-^ secretion and subsequent Cl ^-^/HCO3 ^-^ exchange. At the same time, pH neutral acinar secretion remains undisrupted. This, in combination with reduced CLDN2-dependent pancreatic ductal water secretion, may result in reduced secretory volume with neutral pH and increased digestive enzyme concentration. This would lead to elevated spontaneous pancreatic enzyme activation that could cause pancreatic ductal damage. In addition, in severe cases, thickened pancreatic fluid due to reduced CLDN2 function may cause ductal obstruction, which may contribute to pancreatitis severity. This leads to increased immune cell infiltration and fibrosis. Mechanistically this is analogous to CFTR mutation associated pancreatitis, where the driving force for Na ^+^ and water secretion is reduced by lack of stimulated Cl ^−^ secretion (Angyal *et al*., 2021; Madácsy *et al*., 2018).

Knowledge gained in this study may be used to predict how reported risk alleles may impact pancreatitis development: these polymorphisms may cause reduced functional CLDN2 channels at the tight junction, causing increased sensitivity to pancreatitis. Because these risk alleles are located outside the *CLDN2* coding region, it is possible such reduced CLDN2 function may be caused by decreased CLDN2 expression. It is also possible that additional polymorphisms that are tightly linked with risk allele are responsible for the increased risk of pancreatitis, and such changes could limit CLDN2 expression, channel activity, or channel localization. These possibilities need to be clarified by deep sequencing at the *CLDN2* locus and adjacent region, and functional studies testing how these polymorphisms may affect CLDN2 expression, localization, and function.

CLDN2 risk alleles may intersect with other pancreatitis associated genes. It is reported that *Cftr* KO mice have increased CLDN2 expression in the small intestine, which may compensate for the diminished ion transport (De Lisle, 2014). This points to a possibility that CLDN2 upregulation is also higher in patients with pancreatitis risk loci. It is also possible a fraction of patients with *CLDN2* risk alleles also have other pancreatitis risk alleles, giving them significantly higher risk for pancreatitis and require careful monitoring and more aggressive clinical management. This highlights the importance of genetic testing of *CLDN2* risk alleles.

Taken together, our findings suggest that CLDN2 plays a pivotal role in modulating pancreatic ductal epithelial transport, especially in the context of inflammation, and that its upregulation serves as an adaptive mechanism to mitigate pancreatic injury. These results point to a potential therapeutic target in CLDN2, where activating CLDN2-mediated pathways may promote pancreatic secretion, limit inflammation, and reduce the severity of pancreatitis. Further investigation into the complex interplay between CLDN2, CFTR, and other ion channels could pave the way for novel therapeutic strategies targeting pancreatic ductal function in the management of pancreatitis.

## METHODS

### Human tissue samples

Archival, formalin fixed, paraffin embedded samples of pancreatic tissue were obtained from The Department of Pathology under a protocol approved by The University of Chicago Institutional Review Board. All specimens were from surgical procedures and included six samples of histologically normal pancreas and thirteen samples from patients with non-tumor associated chronic pancreatitis.

### Animals, caerulein-induced experimental pancreatitis, and IFNγ injection

*Cldn2* knockout (KO) mice (*Cldn2*^tm1Lex^) on C57BL/6 background were obtained from Alan S. L. Yu (Kansas University Medical Center, Kansas City, KS)(Pei et al., 2016). Mice were maintained by crossing *Cldn2*^KO/+^ female mice with *Cldn2*^KO/0^ male mice at specific pathogen free barrier facility at the University of Chicago, and all procedures were performed with IACUC approval. Due to *Cldn2* being an X-linked gene, *Cldn2*^KO/+^ female mice and *Cldn2*^+/0^ male mice were used as controls, and *Cldn2*^KO/KO^ female mice and *Cldn2*^KO/0^ male mice were used as the *Cldn2* KO group. This breeding scheme allowed us to circumvent potential CLDN2 dosage effect between male and female control mice. For studies using genetically modified mice, littermates with roughly same numbers of both sexes of mice were used for the study. For caerulein-induced experimental pancreatitis, were subjected to 8 hourly intraperitoneal caerulein injection for two consecutive days (from 9 am each day) at a dose of 75 μg/kg body weight (Moore *et al*., 2019). To determine the early phase of pancreatitis development, pancreatic samples were harvested 2 d after initiation of caerulein injection (1 d after last injection). For the late phase of pancreatitis development, tissue samples were collected 8 d after initiation of caerulein treatment (7 d after last injection) (Moore *et al*., 2019). In a set of experiment, C57BL6/J and *Ifng* KO (B6.129S7-*Ifng^tm1Ts^*/J, JAX #002287) mice was used for caerulein injection.

To determine IFNγ (PeproTech, 315-05) affected CLDN2 expression, C57BL6/J mice were either intraperitoneally injected with PBS, 1×10^4^ U IFNγ dissolved in PBS, or 5×10^4^ U IFNγ dissolved in PBS. Tissue samples were collected 1 day after injection, fixed immediately in 10% neutral buffered formalin for 24 hours, and then transferred to 70% ethanol before undergoing routine paraffin embedding.

### H&E and trichrome staining

Hematoxylin and eosin (H&E) staining of human samples was performed using a Tissue-Tek Prisma automated slide stainer (Sakura) with attached glass cover slipper. H&E staining of mouse pancreas tissue sections was performed manually. Tissue sections were deparaffinized, rehydrated and stained with hematoxylin. After several washes with water and Scott’s bluing reagent, slides were stained with eosin, dehydrated, and mounted. Trichrome staining was performed on formalin fixed paraffin embedded human pancreas tissue and mouse pancreas tissue sections at the Human Tissue Resources Center at The University of Chicago.

### Immunohistochemistry and immunofluorescence staining

Formalin-fixed, paraffin-embedded sections of human pancreas, mouse pancreas, and mouse pancreas organoids were used for immunohistochemical (IHC) and immunofluorescence (IF) staining to detect CLDN2, OCLN, CLDN1, CFTR, and CD45.

For IHC staining of human pancreas sections, slides were deparaffinized, rehydrated, and underwent antigen retrieval by boiling in antigen retrieval solution (Vector Laboratories, Burlingame, CA, USA) for 45 minutes. After washing with 3% hydrogen peroxide, sections were blocked with M.O.M. IgG blocking solution (Vector Laboratories) for 1 hour, rinsed in 2.5% normal horse serum, and incubated for 30 minutes with either anti-CLDN2 (mouse, 1:100, Invitrogen, Cat# 32-5600, RRID: AB_2533085) or anti-OCLN (mouse, 1:100, Invitrogen, Cat# 33-1500, RRID: AB_2533101). Detection was performed using Impress polymer reagent and peroxidase solution (Vector Laboratories), and slides were counterstained with hematoxylin, dehydrated, and mounted.

For IHC staining of mouse pancreas tissue, sections were prepared as above and then incubated with either anti-CLDN2 (rabbit, 1:100, Cell Signaling Technology, Cat# 28530, RRID: AB_2798961) or anti-CD45 (rat, 1:100, Invitrogen, Cat# 14-0451-82, RRID: AB_467251). Staining was conducted at the Human Tissue Resources Center at The University of Chicago.

For IF staining, formalin-fixed, paraffin-embedded sections of human pancreas and mouse pancreas organoids were processed for CLDN2, CLDN1, and CFTR detection. After deparaffinization, sections were washed in 1X TBS-T, quenched with 50 mM NH₄Cl in 1X TBS, and blocked. Sections were incubated overnight with primary antibodies: anti-CLDN2 (mouse, 1:100, Invitrogen, Cat# 32-5600, RRID: AB_2533085) for human tissues, anti-CLDN1 (mouse, 1:100, Invitrogen, Cat# 51-9000, RRID: AB_2533916), or anti-CFTR (mouse, 1:50, Invitrogen, Cat# MA1-935, RRID: AB_2081230). The following day, sections were washed with 1X TBS and stained with Alexa Fluor 594-conjugated donkey anti-mouse secondary antibody (1:1000, Jackson ImmunoResearch, RRID: AB_2340857) and counterstained with Hoechst 33342 (Thermo Fisher Scientific) for nuclear visualization. Slides were mounted with ProLong Diamond antifade mountant (Thermo Fisher Scientific) and imaged on a Keyence BZ-X810 microscope (Keyence Corporation of America, Itasca, IL).

### Quantification of histochemical staining

Trichome, CLDN2, and CD45 stained slides were scanned at 40x magnification using Aperio AT2 slide scanner (Leica Biosystems) and accessed using Aperio eSlide Manager which links to Aperio Imagescope (Leica, v. 12.4.5) software (Leica) for analysis. Slides were annotated manually using Imagescope to select multiple regions of interest for every sample analyzed, which were averaged to provide a single value from each slide for statistical analysis. Trichrome blue staining was quantified using the positive pixel count algorithm v. 8.1 tuned to blue color (Hue center 0.66, Hue width 0.35; Color saturation threshold 0.04, weak intensity 175-220, medium intensity 100-175, strong intensity 0-100). Staining was normalized to total pixels within annotated pancreatic parenchyma. CLDN2 staining was quantified using the positive pixel count algorithm v. 8.1 tuned to brown color (Hue center 0.1, Hue width 0.5; Color saturation threshold 0.04; weak intensity 175-220, medium intensity 100-175, strong intensity 0-100) and normalized to total pixels within the annotated ducts. CD45 positive cells were quantified using automatic cell membrane detection algorithm v. 9.1, with an intensity threshold range of 0 to 220. Lymphoid aggregates and lymph nodes were excluded from analysis. CD45 positive cells were normalized to total cells within the annotation regions.

### Generation and maintenance of organoids

Pancreatic ductal epithelial organoid lines were generated from 6 w old littermate control and *Cldn2* KO mice using published methods, with slight modifications (Huch et al., 2013). Mice were euthanized and pancreas was removed using sterile technique. Pancreas was minced using surgical scissors and digested with 62.5 µg/mL Collagenase from *Clostridium histolyticum* (Sigma, C9407) and 62.5 µg/mL Dispase II (Gibco, 17105-041), 10 µM Y-27632 (Cayman Chemicals, 10005583) and 100 µg/mL Primocin (InvivoGen, ant-pm-1) in Advanced DMEM/F12 medium (Gibco, 12634010), incubating at 37℃ in a shaking incubator for 20 minutes, followed by digestion with TrypLE Express (Gibco, 12605010) and DNase I 0.1mg/mL (Invitrogen, 18047019) for 10 minutes. After trituration, digested samples were filtered through a 100 µm cell strainer. Cells were resuspended in Cultrex UltiMatrix reduced growth factor basement membrane extract (Bio-techne, BME001-05) and plated into prewarmed 24 well plates, and incubated in complete pancreatic organoid growth media (Advanced DMEM/F-12 containing 15 mM HEPES pH7.4 [Gibco 15630130], 50U/mL penicillin-streptomycin [Gibco, 15070063], 100 µg/mL Primocin, 1x serum free B-27 supplement [Gibco, 17504044], 5 mM nicotinamide [MilliporeSigma, N0636], 1.25 mM N-acetylcysteine, [MilliporeSigma, A0737], 10 mM Y-27632, 50 ng/ml EGF (Peprotech AF-100-15), 100 ng/ml FGF-10 [Peprotech, 100-26], 2% Noggin-Fc fusion protein conditioned medium [ImmunoPrecise Antibodies, N002] 2% Rspo3-Fc fusion protein conditioned medium [ImmunoPrecise Antibodies, R001], 5 μM A83-01 [Tocris, 2939], 10 nM [Leu15]-Gastrin I [MilliporeSigma, G9145], 3 μM prostaglandin E_2_ [MilliporeSigma, P0409]). Media was replaced every 2 to 3 d. Pancreatic ductal epithelial organoids were passaged by TrypLE express digestion and split at a 1:3 dilution about every 3-5 days.

For FFPE embedding of organoids, organoids embedded in Cultes UltiMatrix were washed with PBS and freed from extracellular matrix by incubating with cold cell recovery solution (Corning 354253). Organoid pellets were washed with PBS, fixed in 1 ml of 10% neutral buffered formalin, and kept on ice for 1 hour. The fixed organoids were then embedded in HistoGel (Epredia HG4000012), transferred to 70% ethanol, then processed for paraffin embedding and subsequent immunohistochemical analysis.

### RNA Isolation and Real-Time PCR

Pancreatic ductal epithelial organoids were passaged and cultured in differentiation media (complete pancreatic organoid growth media without A83-01, [Leu15]-Gastrin I, and prostaglandin E2) for 2-3 days. Culturing was conducted using suspension culture in low-adherence 6-well plates, with differentiation media supplemented with 2% Cultrex UltiMatrix. After this initial period, the organoid suspension was treated with recombinant cytokines: human IL6 (10 ng/ml, PeproTech, 200-06), murine IL13 (2 ng/ml, PeproTech, 210-13), murine IL22 (2 ng/ml, PeproTech, 210-22), or murine IFNγ (5-20 ng/ml, PeproTech, 315-05) for 2 days. Following treatment, the organoids were spun down, washed with PBS, and total RNA was isolated using the RNeasy Mini Kit (Qiagen, 74104). RNA was reverse transcribed to cDNA according the ProtoScript® II First Strand cDNA Synthesis Kit (New England Biolabs, E6560S). Quantitative PCR was performed with an iQ SYBR Green Supermix (Bio-Rad, 1708882), according to the manufacturers’ instructions and with a Bio-Rad CFX Connect RealTime PCR System. The following primers were used: m*Cldn2*, 5′-GGCTGTTAGGCACATCCA-3′ and 5′-TGGCACCAACATAGGAACTC-3′; mGADPH, 5′ -CTTCACCACCATGGAGAAGGC-3′ and 5′-GGCATGGACTGTGGTCATGAG-3′.

### Time-lapse live imaging of organoids and image quantification

For time-lapse live organoid imaging, control and *Cldn2* KO pancreatic ductal epithelial organoids were digested by TrypLE express then filtered through 40 µm cell strainer. Near single cell suspension was split at 1:100 and embedded in Cultrex UltiMatrix reduced growth factor basement membrane extract in 48 well plates. Organoids were grown in differentiation media (complete pancreatic organoid growth media without A83-01, [Leu15]-Gastrin I, and prostaglandin E2) for 4-5 d until experimentation, with media change every 2-3 d.

To assess changes in organoid size under different conditions, pancreatic ductal epithelial organoids from both control and *Cldn2* KO mice were grown in differentiation media. Organoids were washed with Hanks’ Balanced Salt Solution (HBSS) and pre-incubated in HBSS for 30 minutes. For the Na^+^-gradient dependent paracellular water flux assay, half of the wells for each genotype were switched to regular Na^+^-containing HBSS (mock treatment), while the other half were switched to iso-osmotic HBSS with Na^+^ completely substituted with NMDG^+^. In separate set of experiments to study pancreatic ductal epithelial secretion, after switching to HBSS, organoids were treated with forskolin (10 μM) in the presence or absence of CFTR Inhibitor-172 (20 μM) in HBSS.

Imaging was performed every 10 minutes for 120 minutes using a BZ-X810 Microscope (Keyence) with a 4x Plan Fluorite objective (0.13 NA PhL, Keyence) at ambient temperature to monitor changes in organoid morphology and size. For quantification of organoid size, the acquired images were processed in ImageJ and assembled into time-lapse stacks. The in-house developed OrganoID package(Matthews et al., 2022) was used to measure the area of each organoid, and the data were outputted as Excel (Microsoft, Redwood, WA) documents. Organoid areas at different time points were normalized to their respective starting values for subsequent statistical analyses.

### Data analysis and statistics

Statistical analyses and data visualization was done by using Prism 8.0 (GraphPad, San Diego, CA, USA) and data is reported as mean ± SEM. Two-tailed Student’s t test was used to compare the mean of two groups, one-way ANOVA with Bonferroni *post hoc* analysis was used for multiple comparisons and repeated measures two-way ANOVA analyses was used to compare multiple groups over multiple time points. Statistical significance is designated as *p<0.05, **p<0.01, and ***p<0.001.

## ACKNOWLEDGEMENTS

Organoids were developed by The University of Chicago Organoid and Primary Culture Research Core. Histology and digital pathology services were provided by The University of Chicago Human Tissue Research Center Core. Alan Yu at Kidney Institute, University of Kansas Medical Center kindly provided *Cldn2* KO mice. Schematics were generated using Biorender software. Savas Tay (Pritzker School of Molecular Engineering, The University of Chicago, Chicago, IL, USA) provided analytical tools and conceptual insights. We are grateful for financial support of this work from the following: NIH R01EY027810 (S.A.O.), NIH R01CA219815 (S.A.O.), U01DK127786 (S.A.O.).

## COMPETING INTERESTS

S.A.O is a co-founder, equity holder, and consultant for OptiKIRA, LLC. L.Sh. is a cofounder and equity holder of Nanomics Biotech. C.R.W, L.Sh. and F.K-H. are cofounders and equity holders of Claudyn Biotech.

## AUTHOR CONTRIBUTIONS

S.A.O, L.Sh., and C.R.W. conceived, developed, and supervised the study. S.K., Y.L, J.X., M.T., K.W., A.H., A.K., J.K., N. R., L.Sm., L.Sh., and C.R.W. performed experiments, S.K., Y.L., M.S., S.H., L. Sh., and C.R.W. performed data analysis, M.R. collected clinical data. J.M. and S.T. developed and improved analytical tools, F.K-H. provided analytical tools and conceptual insights. S.K. Y.L., L.Sh., and C.R.W. drafted and revised the manuscript. All authors reviewed and approved the manuscript.

---

**Supplementary Figure 1.**
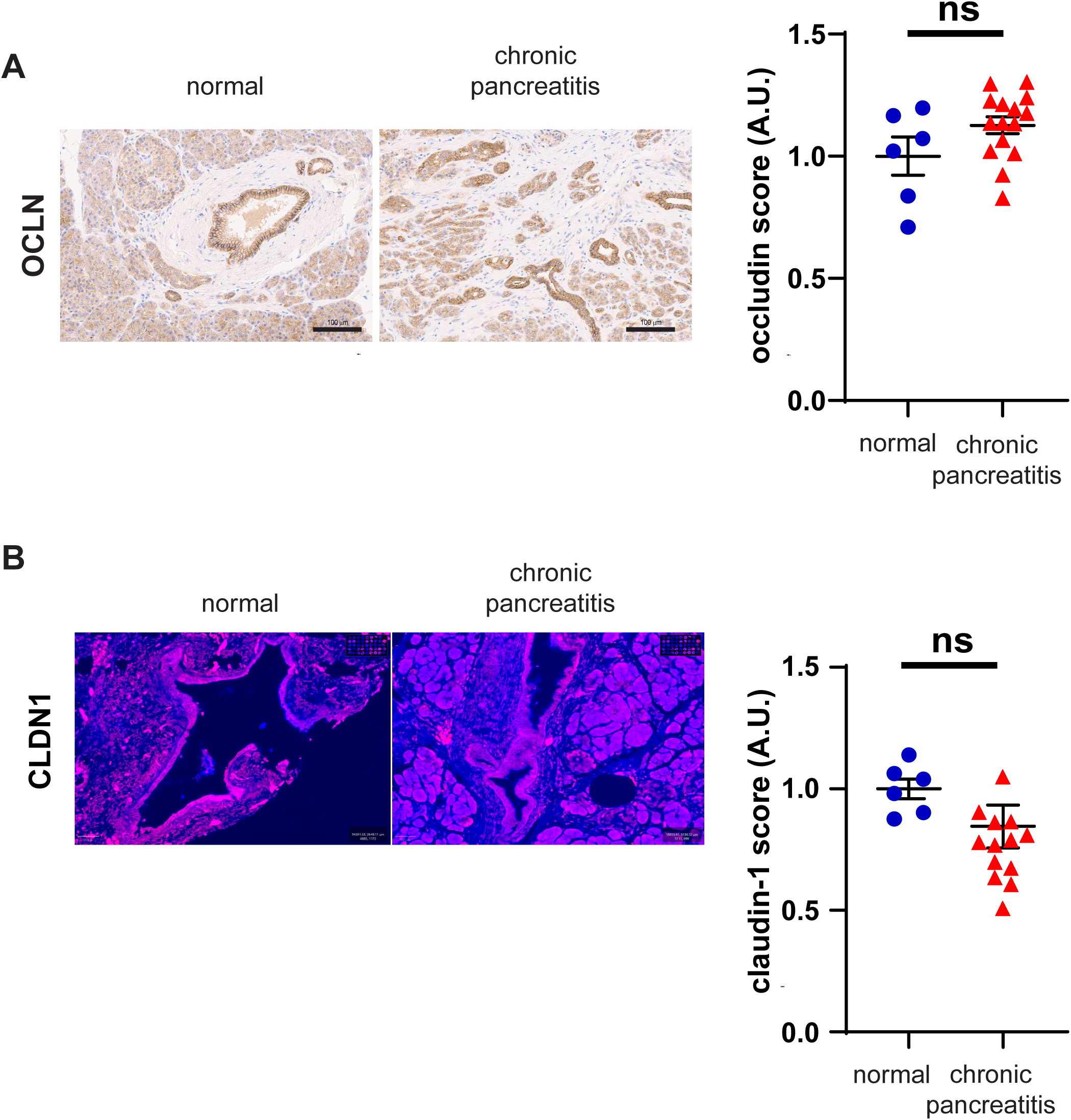
Occludin (OCLN) and Claudin-1 (CLDN1) expression remain unchanged in chronic pancreatitis. **(A-B)** Immunohistochemistry analysis shows no significant difference in OCLN and CLDN1 expression in pancreatic ductal epithelial cells from patients with chronic pancreatitis compared to non-diseased pancreatic tissue (p > 0.05; n = 6-14). Scale bar, 100 µm.

**Supplementary Figure 2.**
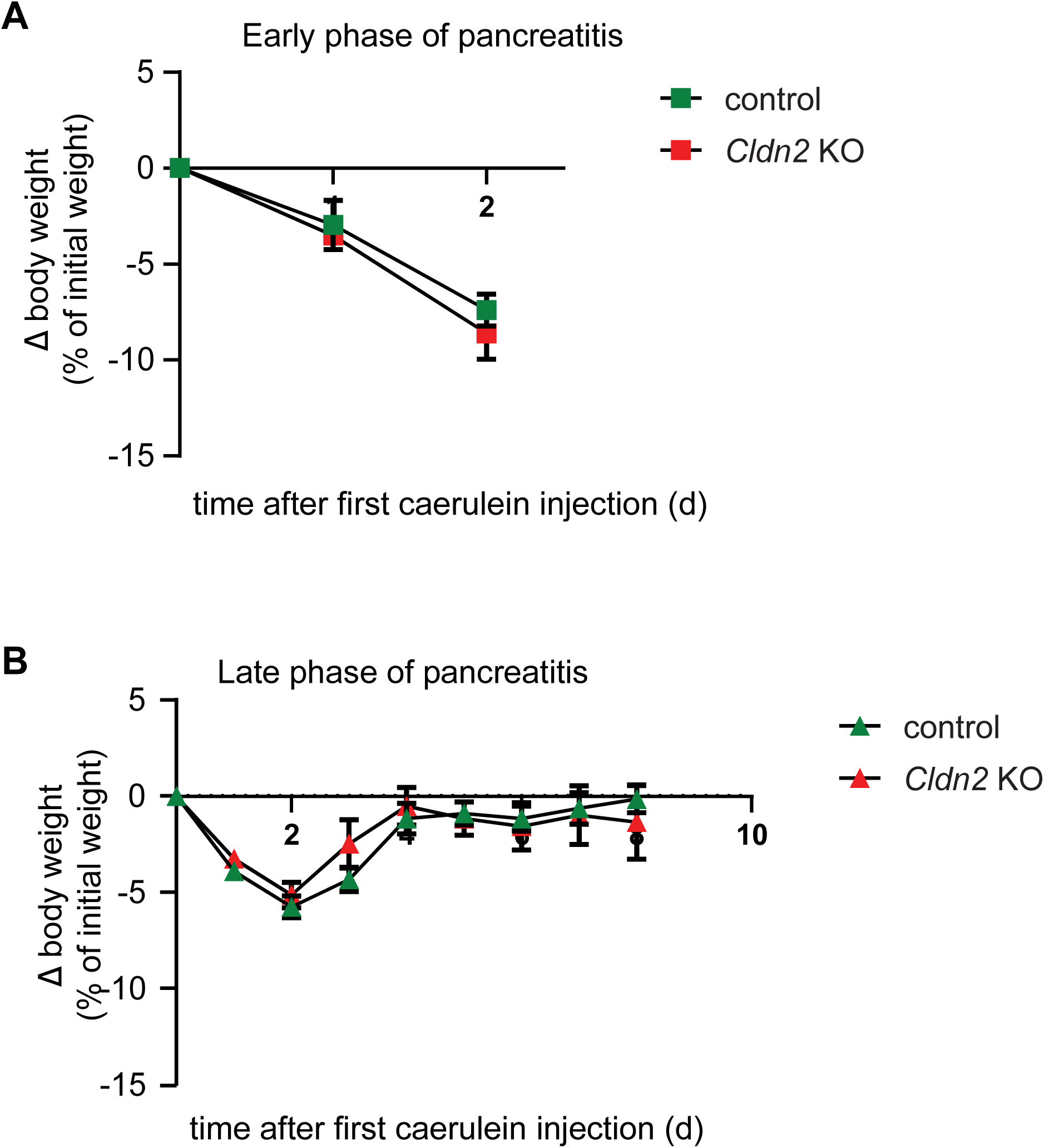
Weight change in mice with caerulein induced experimental pancreatitis. **A.** Body weight change up to 2 d after initiation of caerulein treatment. Green squares and line: caerulein treated control mice (n=4), red squares and line: caerulein treated *Cldn2* KO mice (n=4) (p=0.72). **B.** Body weight change up to 8 d after initiation of caerulein treatment. Green triangles and line: control mice (n=14), red triangles: *Cldn2* KO mice (n=20) (p=0.74). Repeated measures two-way ANOVA was used for statistical analysis.

**Supplementary Figure 3.**
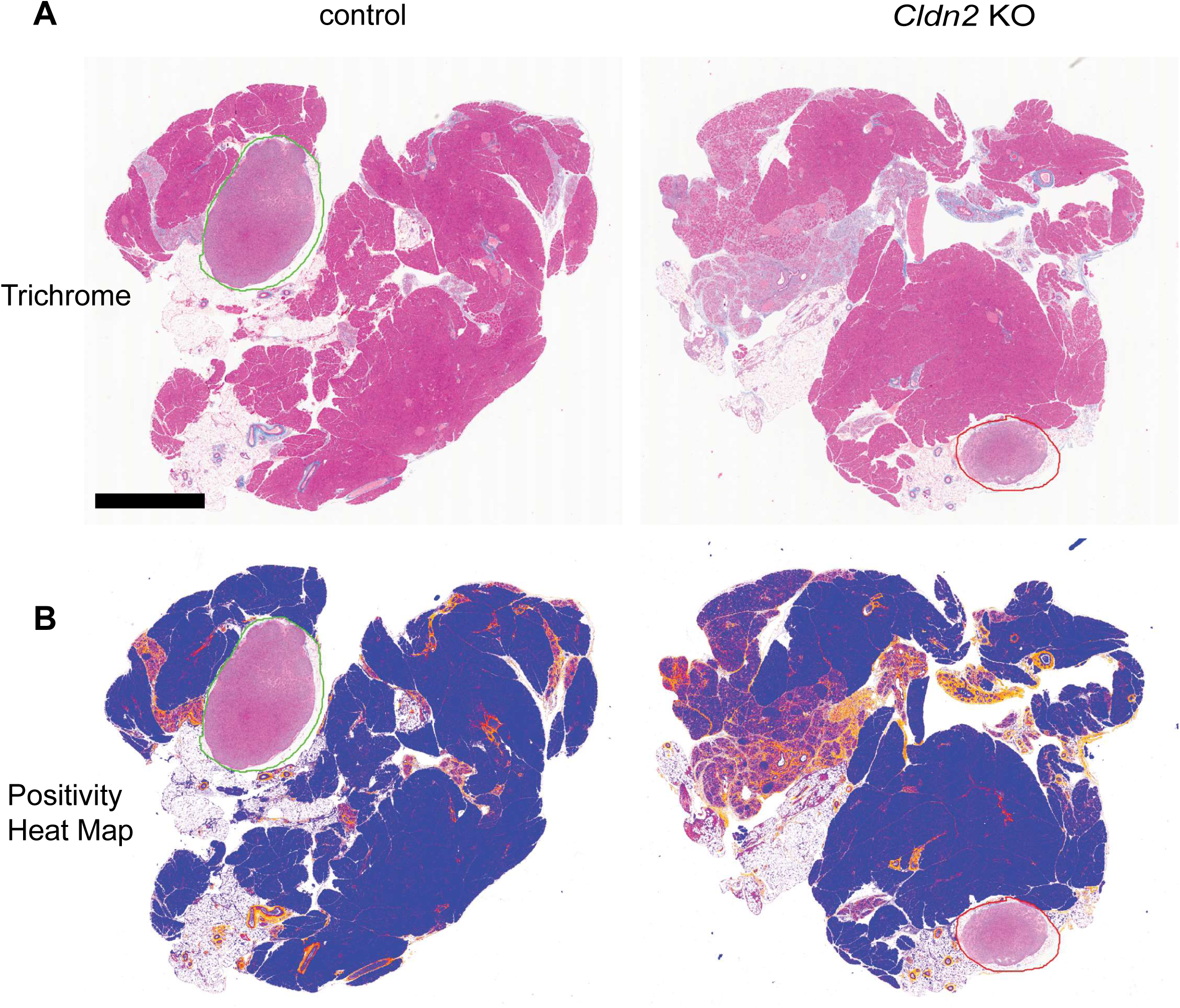
Trichrome stain in 7-day recovery mouse model of pancreatitis. **A.** Trichrome staining of mouse pancreatic tissue sections. (Scale bar 2 mm). Blue staining highlights collagen. The green and red annotations reflect intrapancreatic lymph nodes which were excluded from the trichrome staining analysis. **B**. Heat map of trichrome collagen staining as generated by Aperio Imagescope positive pixel count algorithm. Blue indicates no fibrosis, and yellow, orange, and red indicate increasing degrees of fibrosis. Percent positivity was determined by dividing all positive pixels by total non-white pixels.

**Supplementary Figure 4.**
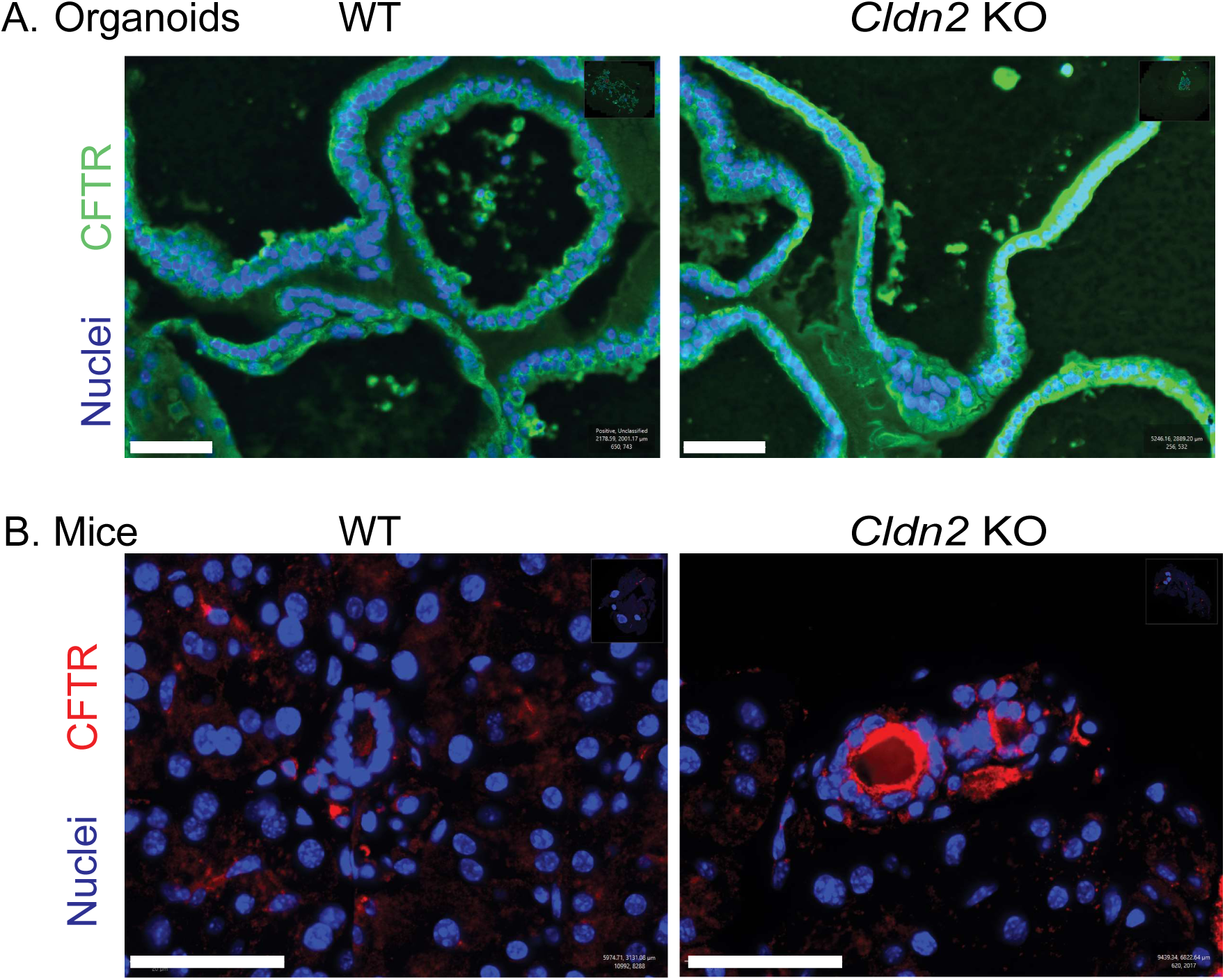
Immunofluorescence staining shows that CFTR expression is maintained in *Cldn2* knockout pancreas organoids and tissue. Scale bar, 50 µm. (A) Pancreas organoids. (B) Pancreas tissue.

## Notes

### Summary of Updates

We have increased the mechanistic understanding behind claudin-2 enhancement in pancreatitis through the following new key findings. 1. Role of inflammation in claudin-2 regulation: We have shown that interferon-gamma drives the upregulation of claudin-2 both in vitro and in vivo, suggesting that pancreatic inflammation can induce claudin-2 expression to limit the development of pancreatitis. This finding provides a novel link between inflammatory signaling and epithelial barrier regulation in the pancreas. 2. Mechanistic Insights into CFTR function in pancreatic ducts: We have demonstrated mechanistically in organoid studies how CFTR is functionally tied to sodium-dependent water transport in pancreatic ductal epithelium. Similar to CFTR inhibition, claudin-2 knockout can block forskolin-mediated secretion in pancreatic ductal organoids. This highlights the integral role of claudin-2 in regulating fluid transport and maintaining pancreatic ductal function under pathological conditions.

